# Mechanical forces at the kidney filtration barrier govern spatial orientation of podocyte processes on capillaries

**DOI:** 10.1101/2023.02.10.528006

**Authors:** David Unnersjö-Jess, Amer Ramdedovic, Linus Butt, Ingo Plagmann, Martin Höhne, Agnes Hackl, Hans Blom, Bernhard Schermer, Thomas Benzing

## Abstract

Mammalian kidneys filter enormous volumes of water and small solutes, a filtration driven by the very high hydrostatic pressure in glomerular capillaries. Interdigitating cellular processes of podocytes form the slits for fluid filtration. They are connected by the membrane-like slit diaphragm cell junction containing a mechanosensitive ion channel complex and allow filtration while counteracting hydrostatic pressure. Using high-resolution microscopy, we show that filtration-slit-generating secondary processes preferentially align along the capillaries’ longitudinal axis while primary processes are preferably perpendicular to the longitudinal direction. The preferential orientation requires maturation in development and is lost in disease states. We demonstrate that loss of proper orientation might contribute to impaired filtration by collapsing of the filtration slits and reducing the mechanical stability of podocyte processes. Together, these data suggest that podocytes sense mechanical strain to utilize circumferential hoop stress balancing the massive mechanical strain generated from fluid flow over the filtration slit.

## INTRODUCTION

The human kidneys produce as much as 180 liters of glomerular filtrate per day in healthy adults, yet only tiny amounts of plasma proteins such as albumin leak into the urine, the end product, with its much smaller volume^1^. The kidney’s filtration barrier is located within microvascular units called glomeruli. It consists of three layers: a capillary endothelium with open pores, a glomerular basement membrane (GBM) of unique composition, and the layer of interdigitating secondary foot processes (FP) of podocytes which are connected by a specialized membrane-like cell junction called the slit diaphragm (SD). Podocytes are post-mitotic cells with limited ability for self-renewal^2^. These cells enwrap the glomerular capillaries and can be lost by detachment from the underlying basement membrane, a culprit in the development of chronic kidney disease^3^. Diseases that reduce the glomerular capillary surface area available for filtration reduce the glomerular filtration rate, affecting the individuals’ overall metabolic balance, leading to detrimental consequences; even small decreases in the glomerular filtration rate are associated with increased cardiovascular morbidity and mortality and reduced overall survival^4,5^.

Although it has been known for many decades that, in the kidney, glomeruli are the site of plasma ultrafiltration and urine production, both the molecular design and function of the filtration barrier remained elusive^1^. However, our recent finding that podocyte FP serve as buttresses against the physical forces of hydrostatic pressure in the glomerular capillaries has provided important insights regarding the physiology of kidney filtration. The buttressing force provided by FP compresses the gel-like structure of the GBM, which in turn restricts the permeability to macromolecules transported by diffusion and bulk flow explaining permselectivity of glomerular filtration^6^. One aspect that was not considered before in our models is that the distending wall stress is not homogeneously distributed on capillaries. Since capillaries to a large degree resemble a cylinder in geometry, the circumferential wall stress will be higher than the axial/longitudinal wall stress. Therefore, we hypothesized that FP might preferentially align along the circumferential axis to maximize the density of buttress-force-generating FP where wall stress, and the consequent need for counteracting forces, is highest.

In addition to wall stress, podocytes are subjected to enormous forces resulting from fluid flow over the filtration slit. Kidney ultrafiltration is driven by the high hydrostatic pressure of about 40 mmHg^1^. Consequently, the filter has to withstand enormous fluid flow. Glomerular filtrate flow represents the highest extravascular fluid flow in the body^7^. Resulting shear stress through the 30 nm-wide slits (with a total slit area of 12,000 μm^2^ per glomerulus) has been calculated to be as high as 8 Pa, acting on the lateral walls of the FP^7^. In addition, fluid flow over the filtration slit will, according to Bernoulli’s principle, lead to a force seeking to pull neighboring FP towards each other. In addition, the hydraulic resistance from the slit diaphragm protein complex will lead to additional forces, acting to collapse the slit diaphragm. It might therefore also be mechanically beneficial to align the SD molecules in parallel with the highest wall stress in order to counteract such forces. Although it has been known for a long time that the slit diaphragm is a non-distensible structure^8^ and that podocyte foot processes contain a sophisticated actin cytoskeletal machinery that is regulated through signaling at the slit diaphragm^9^, the biophysical basis of capillary wall stabilization is still unclear.

The question of whether or not FP display a random orientation on glomerular capillaries, and how a hypothetical preferred orientation of FP might have a role for the function of the kidney filter, has been discussed in previous studies^10,11^. It has however not to date been quantitatively and convincingly described if FP do show a preferred orientation on glomerular capillaries or if the distribution is random. The reason for this is probably that the three-dimensional geometry of glomerular capillaries is rather complex, making it challenging to define the orientation of the convoluted capillary from electron microscopy (EM) data. Further, it is with all modalities of EM challenging to cover the full depth of a glomerular capillary (~ 10 μm), at least at sufficient throughput to quantitatively describe the orientation of large numbers of FP on the curved capillary surface. Recently, we have developed new sample preparation and imaging tools which allow for high-throughput quantitative 3D imaging at a resolution sufficient to resolve FP^6,12–15^. Using these novel tools, we here describe the novel finding that secondary FP preferentially align along the longitudinal axis of the capillaries, an alignment which takes place as the filtration barrier matures in development. This indicates that podocytes appear to respond to the mechanical forces that occur with the development of pressurized vasculature.

## RESULTS

### Podocyte processes show a preferred orientation on glomerular capillaries of wild-type mice

To be able to analyze whether FP display a preferred orientation on capillaries, we assumed that a statistically relevant analysis would require a very large number of FP to be included. Thus, in order to ensure highest data throughput, we applied our latest sample preparation protocols which induce slight swelling of tissue samples (around 30%)^14,15^. This allows for resolving nanoscale FP using confocal microscopy, drastically simplifying image acquisition (as compared to protocols which rely on super-resolved microscopy). With the simplified protocol, glomerular capillaries can be imaged in 3D allowing both for determining capillary orientation as well as providing sufficient spatial resolution for resolving and determining the orientation of FP (Fig. 1 a-c, Supplementary Movie 1). Primary processes (PP) are visualized by staining for acetylated tubulin and FP are visualized by staining for the SD using a nephrin antibody. Importantly, since samples are thick and transparent, the protocols allow for focusing in depth onto suitable regions-of-interest, where glomerular capillaries can be imaged in their full extent (a problematic task in 4-10 μm thin paraffin/frozen sections). To quantitatively assess the orientation of FP with regard to the capillary orientation, we used the strategy described in Fig. 1a-b (see figure legend and methods section for details). We were only interested in evaluating whether FP are oriented in parallel or perpendicularly to the capillary orientation, and thus all angles (calculated with regards to the x-axis from line profiles as shown in Fig. 1b) were projected onto the first quadrant where 0° is parallel and 90° perpendicular with regard to the capillary axis (0°-90°, e.g. the angle 270° → 90°, 180° → 0° etc., see methods section for more details). After determining orientation angles for over 6800 FP in 3 different wild-type mice, we observed a clear and significant preference for foot processes in healthy mice to be oriented in parallel with the capillary orientation (Fig. 1c, left panel). Since foot processes are known to protrude perpendicularly from PP (like branches off a tree), we also analyzed the orientation of PP. Indeed, we found that PP showed a preferred orientation perpendicularly to the capillary orientation, further adding support to our finding and acting as an internal control that the analysis procedure itself does not bias towards any specific orientation (Fig. 1c, right panel). Importantly, the findings in Fig. 1e-f were reproducible between the three WT mice included in the analysis (Supplementary Fig. S1). Thus, these results clearly suggest that foot processes are preferably oriented perpendicularly to the circumferential axis, where wall stress is highest, and consequently a need for counteracting forces is also the highest.

**Figure 1.**
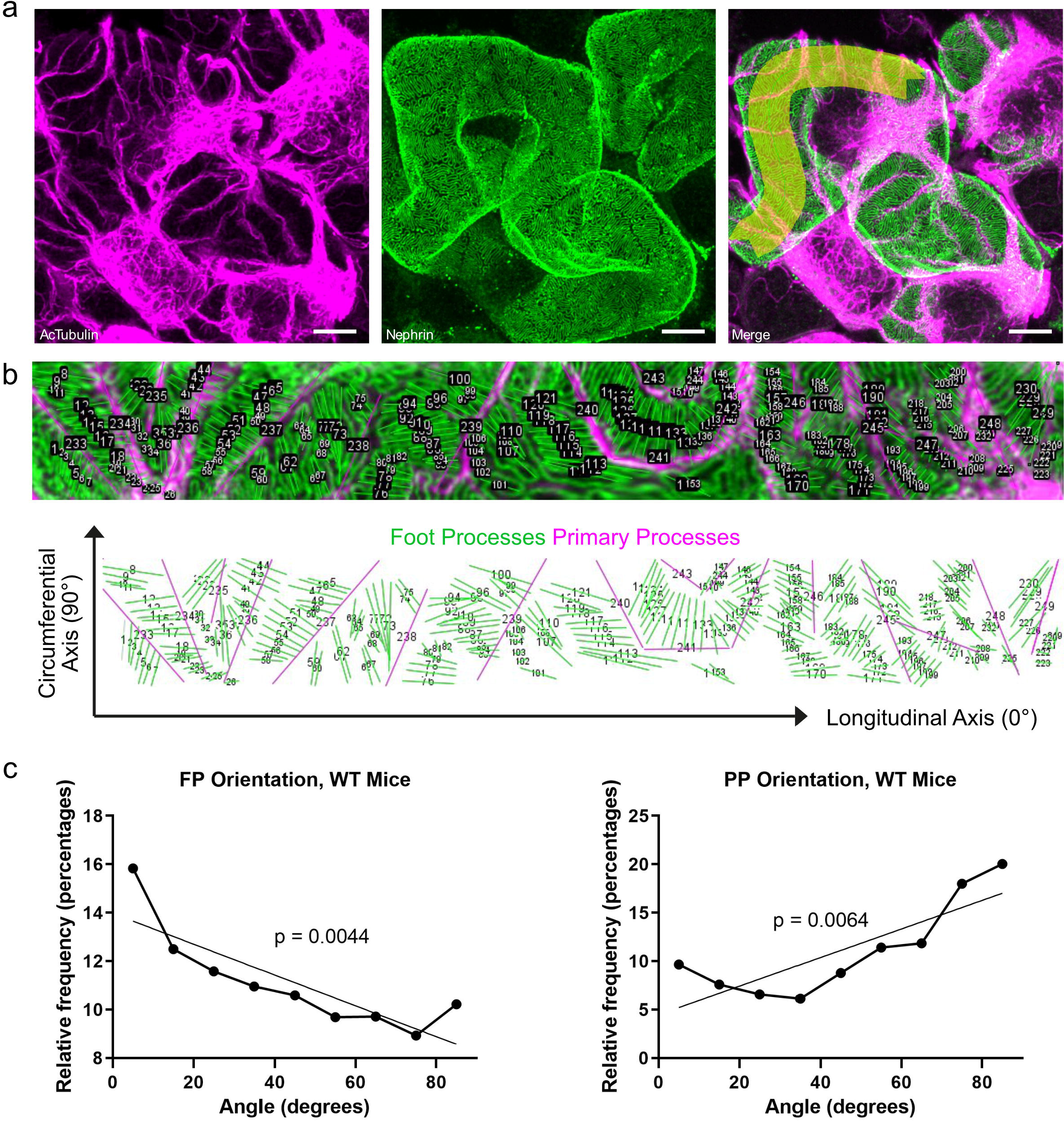
The filtration slits preferentially align along the capillary axis. Deconvolution of three-dimensional optical imaging of glomerular capillaries resolves the angular distribution of podocyte primary (PP) and secondary foot processes (FP), and filtration slits relative to the longitudinal axis of glomerular capillaries. Scale bars 5 μm. **a** Maximum intensity projection of 15 μm thick z-stack stained for acetylated tubulin with Cy3/magenta (primary processes (left panel)) and nephrin with Alexa-488/green (slit diaphragm/foot processes (middle panel)) shows that the entire surface of glomerular capillaries can be fully visualized with high resolution using confocal microscopy. To allow deconvolution of the curved capillary, the orientation of glomerular capillary segments is outlined using the “thick line selection” in ImageJ as indicated in yellow (right panel). **b** Deconvolution of the three-dimensional image using the ImageJ “straighten” plugin. Deconvoluted (straight) capillary projected onto the longitudinal axis. FP (green line selections) and PP (magenta line selections) are annotated in the 2D deconvoluted image (upper panel). The lower panel shows these annotations against a white background with the longitudinal/circumferential axes drawn out. **c** Angular distribution of FP (left panel) and PP (right panel) relative to the x-axis (deconvoluted longitudinal axis of the capillaries) projected onto the first quadrant (0 degrees = parallel with capillary orientation, 90 degrees = perpendicular to capillary orientation). Statistical significance was calculated by fitting a line to the distribution curve using linear regression, and then calculating the p-value for this line to have a non-zero k-value (k-value of 0 corresponds to a straight horizontal line = random distribution). p-values are indicated next to the regression lines in the graphs. There is a clear and significant preference for FP to be oriented in parallel with the capillary orientation (over-representation of angles close to 0 in (**c**, left panel)) and for PP to be oriented perpendicularly to the capillary orientation (over-representation of angles close to 90 degrees in (**c**, right panel)).

### A preferred podocyte process orientation is neither observed in newborn mice nor in glomerular disease mouse models of different origin

We next went on to study the orientation of podocyte processes in a newborn P0 wild-type mouse, where glomerular vascularization and development of podocyte processes is not completed. It is therefore expected that the mean capillary wall pressure needed to be counteracted by foot processes is lower in these mice than in the case of adult mice. Interestingly, in P0 mice, we did not observe the preferred orientation of foot processes as observed for adult mice (see Fig. 1), possibly as a result of lower wall stress but also possibly due to the fact that podocyte development is at this stage not fully completed (Fig. 2a). To study how glomerular disease affects the orientation of foot processes, we additionally performed the orientation analysis in two different mouse disease models; mice injected with nephrotoxic serum (NTS) and mice with two compound-heterozygous mutations in the NPHS2 gene (R231Q/A286V). NTS mice develop glomerulonephritis with mild effacement of FP within 10 days of injection, while R231Q/A286V mice develop an FSGS histological phenotype at around 8 weeks of age with more severe effacement of FP. We observed a loss of podocyte process orientation in both these disease models of different origin (non-genetic/genetic) (Fig. 2b-c). According to our hypothesis, a preferred orientation of FP is for the purpose of aligning SD molecules along the higher circumferential wall stress component. We have previously shown that upon FP effacement, FP take on a more circular shape^6^, which might explain the lack of an optimized orientation in glomerular disease. For a circular FP, there is simply no orientation which will maximize the number of SD molecules aligned in parallel to the circumferential axis (Supplementary Fig. S2).

**Figure 2.**
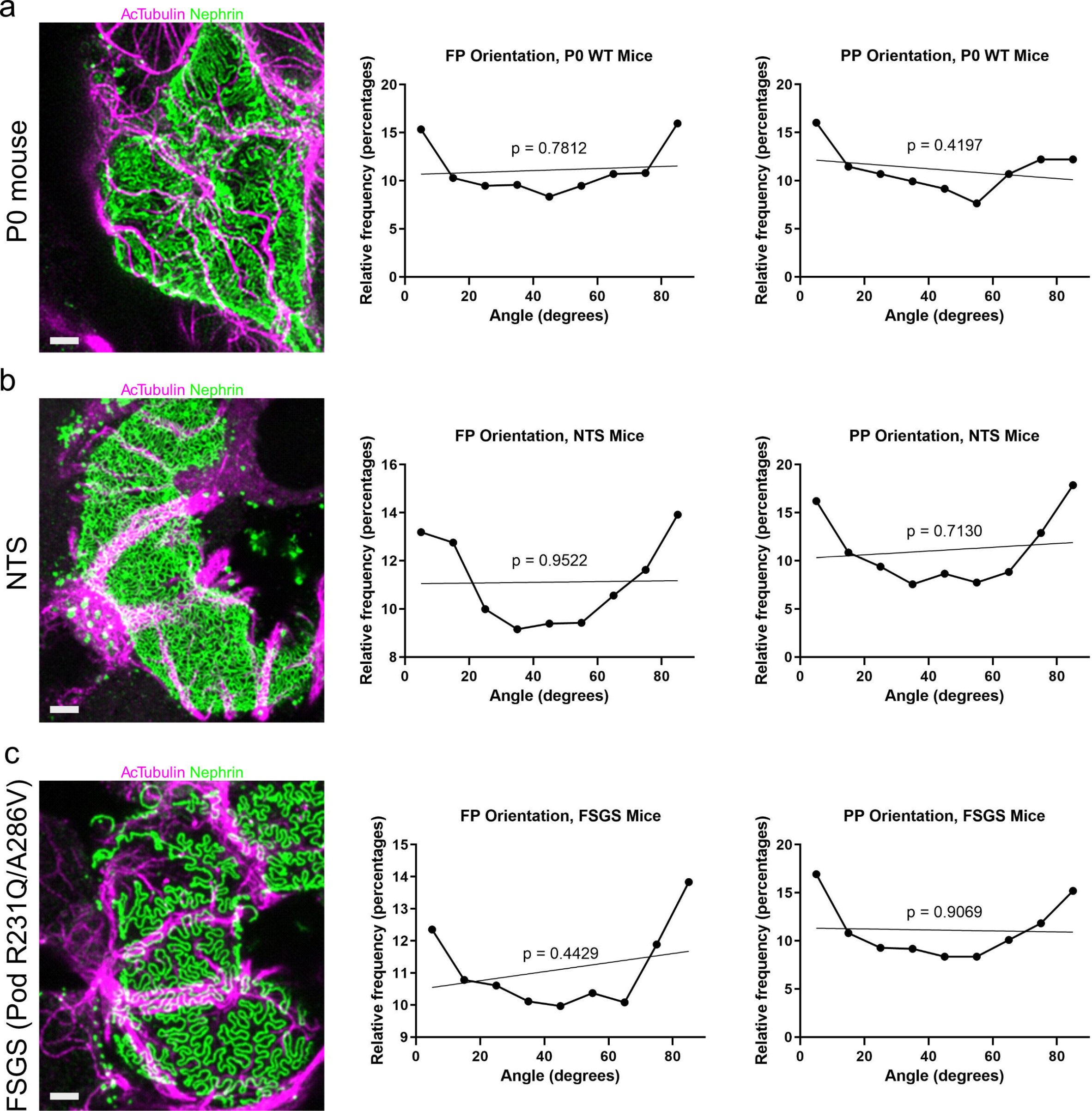
Coordinated FP orientation is lost in early stages of genetic and acquired kidney disease and requires completed development. **a**, Representative image of a glomerular capillary from a P0 wild-type mouse (left panel) along with angular distributions of foot processes (middle panel) and primary processes (right panel) of three independent animals. In contrast to human kidneys, kidneys of newborn mice are not fully developed but require another few days to mature. No significant preference of orientation is observed at P0. **b**, Representative image of a glomerular capillary from a mouse injected with nephrotoxic serum (NTS, left panel) along with angular distribution of foot processes (middle panel) and primary processes (right panel). No significant preference for orientation was observed. **c**, Representative image of a glomerular capillary from a mouse with two compound heterozygous mutations in the *Nphs2* gene (R231Q/A286V) and early disease manifestation, along with angular distribution of foot process (middle panel) and primary process (right panel). No significant preference for orientation is observed. Statistical significance was calculated by fitting a line to the distribution curve using linear regression, and then calculating the p-value for this line to have a non-zero k-value (k-value of 0 corresponds to a straight horizontal line = random distribution). p-values are indicated next to the regression lines in the graphs. Stainings for PP and FP same as in Fig. 1. Scale bars 2 μm.

### A preferred podocyte process orientation is only found in straight, cylinder-like, segments of glomerular capillaries

If FP are truly oriented according to wall stress components, the orientation preference should be more pronounced in long and straight capillary segments, where the resemblance of the capillary to cylindrical geometry is highest. In curved segments, where the capillary makes a sudden turn, the capillary surface will, locally, more resemble a sphere in shape, with a consequently more homogeneous distribution of wall stress on the surface. We therefore went on to study the angular distribution in straight and curved capillary segments (Fig. 3). By separately analyzing straight capillary segments, we found an even more pronounced and significant distribution preference of both FP and PP (Fig. 3a). When analyzing curved segments of glomerular capillaries, we did not observe a preferred orientation (Fig 3b). These results further support our hypothesis that the preferred spatial orientation is a result of mechanical forces acting on podocytes.

**Figure 3.**
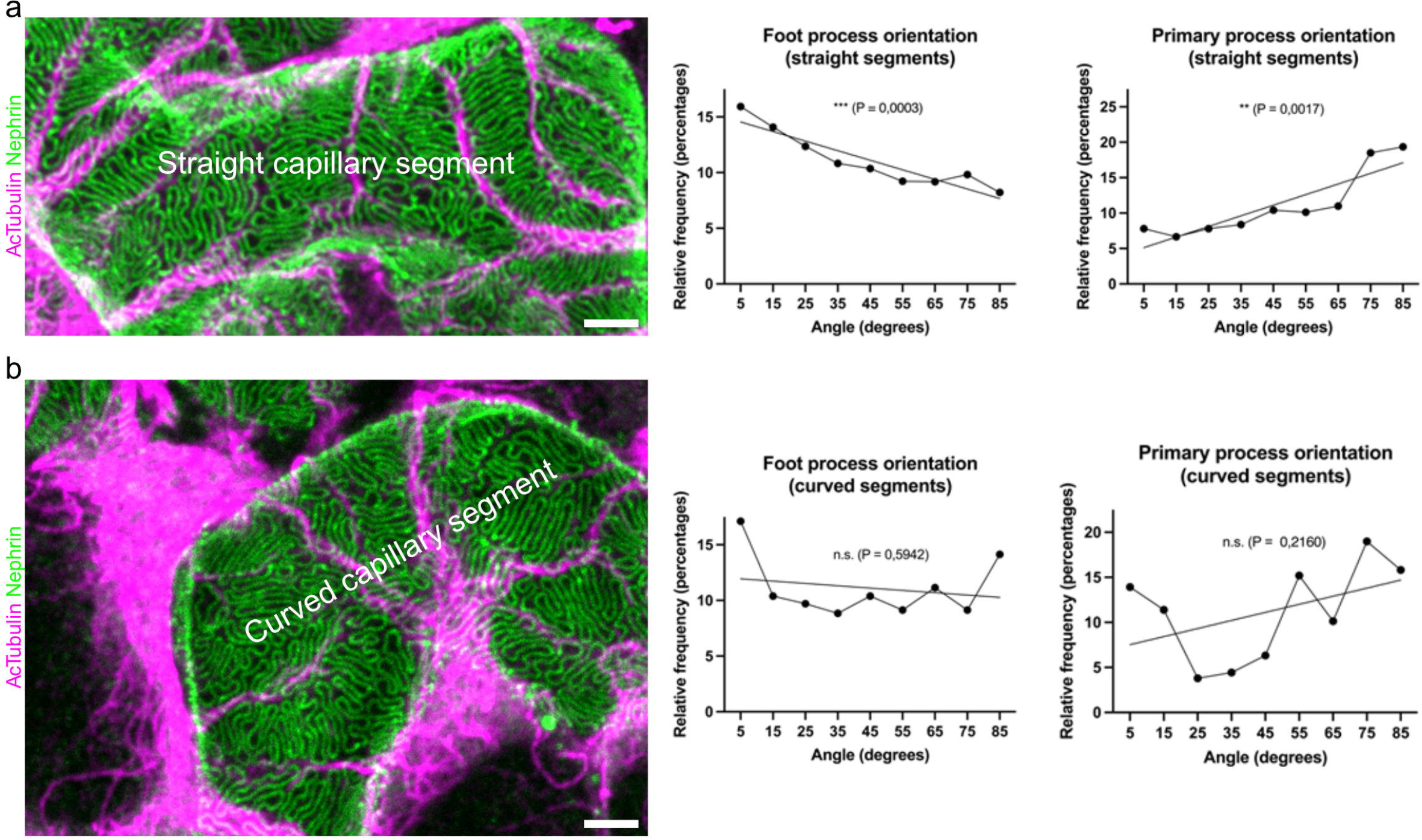
Analysis of straight and curved segments of glomerular capillaries reveals an adaptive (active) orientation of podocyte processes in areas of differences in wall stress. Podocyte process orientation analysis in straight and curved capillary segments. Of the ~ 6800 FP from Figure 1, ~ 2800 were assigned to straight and ~ 1000 to curved capillary segments, respectively. Only obviously straight or curved capillary segments were included. The scale bar is 2 μm. **a**, Representative image of a glomerular capillary with a straight appearance (wild-type animal) along with angular distribution of FP (middle panel) and PP (right panel) in all the included straight segments. **b**, Representative image of a glomerular capillary segment with a curved appearance (left panel) along with angular distribution of FP (middle) and PP (right) in all the included curved segments. The angular distribution is more randomized, with only primary processes showing a slight tendency to be oriented perpendicularly to capillary orientation suggesting an adaptive orientation of podocyte processes in those areas which display differences in circumferential hoop stress and longitudinal forces. Statistical significance was calculated by fitting a line to the distribution curve using linear regression, and then calculating the p-value for this line to have a non-zero k-value (k-value of 0 corresponds to a straight horizontal line = random distribution). p-values are indicated next to the regression lines in the graphs. Stainings for PP and FP same as in Fig. 1.

## DISCUSSION

Recent advances have led to the development of an experimentally validated biophysical model of kidney ultrafiltration^1,6^. However, how podocytes may resist the enormous mechanical strain at the filtration barrier and why the filtration slits do not collapse through massive fluid flow has remained elusive. Addressing these essential questions in kidney physiology has not been possible in the past due to technical limitations. Here, we used ultrahigh-resolution imaging, image analysis, and genetically engineered mouse models of human disease to show that podocyte processes are not randomly organized on glomerular capillaries, but are spatially aligned according to mechanical forces acting on them. This orientation preference is only observed in adult wild-type mice, and is absent both in newborn mice and in two disease models of different origin (genetic FSGS, inflammatory NTS). The fact that the orientation preference is only observed in straight, cylinder-resembling parts of capillaries provides strong support that the orientation results from mechanical forces acting on podocytes.

Assuming capillaries are considered thin-walled cylinders (wall thickness smaller than capillary radius by ~ a factor of 50, a condition fulfilled in glomerular capillaries), radial wall stress is neglectably low as compared to the longitudinal and circumferential wall stress. The longitudinal wall stress component will in turn be of lower magnitude as compared to the circumferential wall stress by a factor of two if the capillary is assumed to have closed ends^16^. If the capillary is assumed to have open ends, there will be no axial wall stress component. The situation in a glomerular capillary is likely somewhere in between these two extreme cases, but it will always hold true that, for cylindrical geometry, the circumferential wall stress component will be significantly larger than the other components (Fig. 4a). With the observed orientation preference, FP will be aligned in parallel with the orientation axis of the capillary. This has both the consequence that the number of FP and the density of SD is higher along the circumferential axis, and also that SD molecules (e.g. nephrin, neph1 etc.) are aligned in parallel with the circumferential axis. It has previously been proposed that the molecular complex of the so-called slit diaphragm (SD), the cell contact between interdigitating FP, exhibit spring-like properties, possibly to react/respond to forces resulting from capillary pressure^17^. It would therefore be beneficial to align these spring-like molecules along the axis where wall stress is highest. It has further been suggested that the SD likely exhibits a substantial hydraulic flow resistance, which might result in a “bending” of SD molecules, with resulting forces acting to decrease the width of the SD^18,19^. Further, according to Bernoulli’s principle, the faster fluid flow in the narrow slit will result in additional forces acting to collapse the slit. Alignment of the SD in parallel with the highest wall stress component might therefore be for the purpose of counteracting these forces in order to keep the junction optimally open to filtration flow (Fig. 4b).

**Figure 4.**
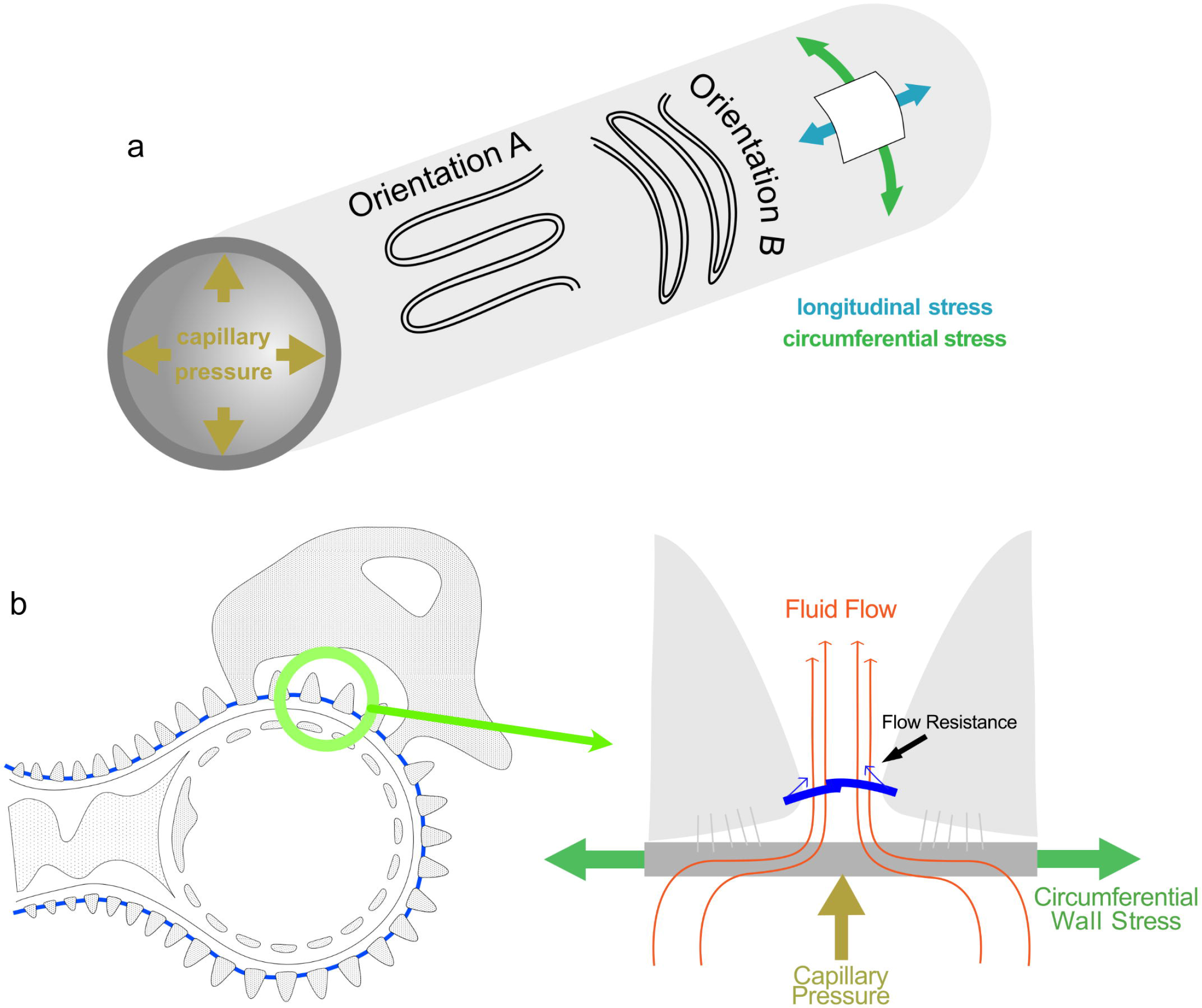
Differences in circumferential stress and longitudinal stress are sensed by the podocyte to preferentially align FP along the capillary axis. Schematic summary of results, with biophysical interpretations. **a**, A simplified schematic drawing of a glomerular capillary, with intraluminal blood pressure (brown/yellow) resulting in circumferential (green) and axial/longitudinal (blue) wall stress. For cylindrical geometry, the circumferential wall stress will always be larger in magnitude than the axial wall stress (in ideal thin-walled closed cylinders a factor of two larger). Depicted are also the two extreme cases of foot process orientations, where orientation A is the observed preferred one. Orientation A will maximize the one-dimensional density of foot processes along the circumferential axis, where the need to counteract wall stress is highest. In addition, orientation A will orient slit diaphragm molecules, such as nephrin, in parallel with the highest wall stress, allowing the spring-like properties of these molecules to help coping with circumferential wall stress. **b**, The parallel orientation of FP might also be beneficial from a hydrodynamics point of view. The left panel shows a capillary loop covered with FP aligned in parallel with the capillary orientation. Right panel shows a zoom in of two neighboring FP, where the fluid flow over the slit diaphragm (yellow/orange arrows) will be hindered by hydraulic resistance at the SD. This, together with Bernoulli’s principle, will cause a bending of SD molecules, leading to forces acting to narrow the filtration slit (blue arrows). The observed preferred orientation of FP will align the SD along the highest wall stress component which will counteract the narrowing of the filtration slit. Therefore, the observed orientation will be advantageous in terms of keeping the SD open to filtration flow.

The lack of orientation preference in two different mouse models for kidney disease can be explained by the fact that. for these effaced, more circular FP, there is no orientation which maximizes the density of SD along the axis of highest wall stress. Thus, the lost orientation of FP in disease is further indication that the orientation preference we observe in this study is for the purpose of aligning SD molecules along the axis where wall stress is highest.

In summary, this study reports the novel discovery that foot processes are not randomly distributed on glomerular capillaries, a subject which has been debated for decades. We here, based on theoretical reasoning, experimental data and previous studies, suggest physical reasons for the observed orientation preference. We have in this study however not addressed and deciphered the presumably complex and multi-factorial molecular mechanism by which a correct orientation of FP is maintained by the podocyte. There is however within the field of mechanobiology emerging evidence that podocytes possess a mechanosensitive machinery located at the slit diaphragm, and that mutations in such mechanosensitive proteins lead to glomerular disease^20^. It is based on this intriguing to speculate that the SD may not only act as a spring-like structure designed to respond to forces on the capillary wall, but also possesses mechanosensory properties in order to react to mechanical cues. These cues might result in the transduction of downstream signals for reorganization of cytoskeletal proteins, such as actin, and cell adhesion proteins, such as integrins, for the purpose of spatially orienting the nanoscale podocyte processes on the capillary wall.

## METHODS

### Mouse and human kidney tissue

All mouse experiments were approved by the State Office of North Rhine-Westphalia, Department of Nature, Environment and Consumer Protection (LANUV NRW, Germany) and were performed in accordance with European, national and institutional guidelines. Mice of 100% C57BL/6N background were used. After anesthesia with Ketamine and Xylazine, mice were euthanized by cardiac perfusion with Hank’s Balanced Salt Solution (HBSS) and fixated as described below.

Control human tissue was collected from patients that were nephrectomized due to renal tumors. Tissue sample was dissected from the non-tumorous pole of the kidney and showed normal histological picture in routine histological examination.

### Mouse Model for FSGS

Mice with two compound-heterozygous point mutations, Pod^R231Q/A286V^, were generated as previously described^6^. Mice were sacrificed at 0 or 4 weeks of age as stated above.

### Nephrotoxic Serum (NTS) Injection

10 days prior to sacrifice, 11 μl/g body weight of a sheep anti-rat whole glomerular antibody (PTX-001S-Ms, Probetex Inc, San Antonio, TX, USA) was injected intraperitoneally in a 3-week-old mouse^13^. The mouse was euthanized and kidneys were dissected and fixated as described above.

### Preparation of kidney tissue for confocal microscopy

Experimental mice were euthanized by decapitation (only newborn mice) or cardiac perfusion with Hanks’ balanced salt solution (HBSS; 5.4 mM KCl, 0.3 mM Na_2_HPO_4_, 0.4 mM KH_2_PO4, 4.2 mM NaHCO_3_, 137 mM NaCl, 5.6 mM D-glucose, 1.3 mM CaCl_2_, 0.5 mM MgCl_2_, 0.6 mM MgSO_4_) following cardiac blood draw and anesthesia with Ketamine (Zoetis) and Xylazine (Bayer). After median laparotomy, kidneys were removed and fixed in 4% neutral buffered formalin for 2-4 hours at room temperature or overnight at 4°C. Kidney tissue was prepared with a recently published fast protocol^14^. Here, fixed kidneys were sectioned using a Vibratome before incubation in clearing solution (200 mM boric acid, 4% SDS, pH 8.5) for 1-2 h at 70°C. Prior to immunolabelling, samples were washed in PBST (0.1% Triton-X in 1X PBS) for 10 min. For formalin-fixed paraffin-embedded (FFPE) tissue, samples were treated according to a recently published protocol optimized for FFPE samples^15^. Small pieces of kidney tissue were cut directly from paraffin blocks, transferred to 1.5 mL Eppendorf tubes, de-paraffinized using xylene and re-hydrated using decreasing concentrations of ethanol in DI water. After this, sections were incubated in clearing solution for 2 hours at 90°C, before proceeding with immunolabelling after washing in PBST as for the other samples.

### Immunolabelling

Samples were incubated with primary antibody diluted in PBST for 24 h at 37°C, followed by washing in PBST for 5 min and after this incubated in secondary antibody diluted in PBST. Samples were finally washed in PBST for 2*5 min. Primary antibodies used: Sheep anti-nephrin (R&D Systems AF4269 1:50), mouse anti-acetylated tubulin (SIGMA T6793 1:100), guinea pig anti-nephrin (Fitzgerald 20R-NP002 1:50). Secondary antibodies used: donkey anti-sheep antibody coupled to Alexa-488 (Jackson ImmunoResearch 713-005-147 1:50), donkey anti-mouse antibody coupled with Cy3 (Jackson ImmunoResearch 715-165-150 1:50), goat anti-guinea pig secondary antibody coupled with Alexa-488 (Abcam ab150185 1:50).

### Mounting

Samples were incubated in 80% wt/wt fructose (1 mL of dH_2_0 added to 4 g of fructose) containing 4 M Urea at 37°C with shaking at 500 rpm for 15 minutes and then placed in an Ibidi dish. To prevent evaporation, a cover slip was placed on top of the sample prior to imaging.

### Imaging

Confocal images were acquired using a Leica SP8 system using a 100X/1.4 NA objective. As previously described, a confocal pinhole setting of 0.3 airy units (AU) (as calculated for 594 nm light) was used to ensure sufficient spatial resolution to resolve FP^14^.

### Orientation Analysis

All analysis was carried out using Fiji^21^ on maximum-intensity-projected z-stacks as the ones shown in Fig. 2–4. As indicated in Fig. 2, the capillary was manually annotated by using a line profile in ImageJ. The thickness of the line profile was adjusted to include most of the capillary surface. After this, the “straighten” plugin in ImageJ was used to project this capillary segment onto the x-axis. In Figure 2, the yellow line profile results in the straightened capillary segment in panel (b). In short, this tool fits a cubic spline to the line selection and straightens the object along it. More information can be found at https://imagej.nih.gov/ij/plugins/straighten.html. After this, foot processes and primary processes are annotated using straight lines as shown in Fig. 2. The line is always drawn from the base of a process to the tip of the same process. Occasionally, as can be seen for a primary process in the middle of panel (d) of Fig. 2, highly bent processes are divided into two or more line-profiles of equal length. After annotating processes, angles were measured with respect to the x-axis using the “measure” tool in ImageJ. This tool returns angles between ± 0-180°. All of these angles are projected on to 0-90°. For negative angles between 0-90, the absolute value was taken, e.g. −35° → 35°. For negative angles between 90°-180°, 180° was added, so that e.g. −135° → 45°. For positive angles between 90°-180°, 180° was subtracted and the absolute value was taken, so that e.g. 135° → 45°. After this, frequency distributions were generated using Graphpad Prism. ~ 6800 FP and ~ 800 PP were analyzed in at least three different mice for each data point.

## Supporting information

3D-rendering of a z-stack showing podocyte processes on glomerular capillaries in a wild-type mouse. Stainings and colors same as Fig. 1.

## ACKNOWLEDGEMENTS

We thank the CECAD Imaging Facility, Cologne, Germany and the Advanced Light Microscopy (ALM) Facility, Solna, Sweden for their support in the acquisition of microscopy data. This work was supported by the Clinical Research Unit of the Deutsche Forschungsgemeinschaft (DFG) CRU 329 to T.B. and B.S. as well as by the Research Unit of the DFG FOR 2743 to D.U.-J. and T.B.. DUJ is partly funded by grants from Njurfonden and Torsten Söderbergs Stiftelse. A.H. is a holder of a stipend from the Peter Stiftung.

## AUTHOR CONTRIBUTIONS

Conceptualization, B.S., T.B, H.B., D.U-J.; methodology, D.U-J, A.R., L.B., I.P., M.H., H.B., A.H., B.S. and T.B.; data analysis, D.U-J., A.R., investigation, D.U.-J., A.R., L.B., I.P., M.H.; writing, original draft, D.U-J., T.B.; writing, review and editing, D.U-J., A.R., L.B., I.P., M.H., H.B., A.H., B.S., T.B.

## DATA AVAILABILITY

The data which supported the findings in this paper can be provided by the corresponding authors upon request.

