## Supplementary Figures for "Mechanical forces at the kidney filtration barrier govern spatial orientation of podocyte processes on capillaries"

**Running title:** Role of foot process orientation in albumin filtration

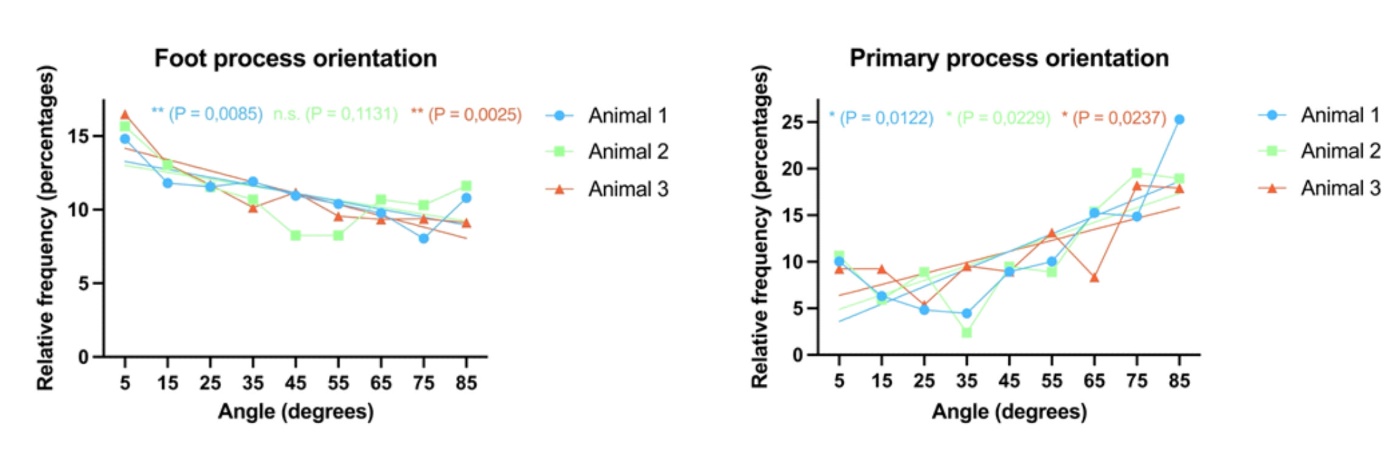

**Supplementary Figure 1.** Reproducibility of preferred orientation between individual animals. Frequency distributions of foot process (left) and primary process (right) for three different WT mice (color coded). All animals show a significant orientation preference for both primary processes and foot processes, with the exception of foot processes for animal 2 in the left panel, however, the tendency is for this animal clearly towards lower angles with a p-value of 0.1131. Statistical significance was calculated by fitting a line to the distribution curve using linear regression, and then calculating the p-value for this line to have a non-zero k-value (corresponding to a straight horizontal line = random distribution). p-values are indicated next to the line in the graph.

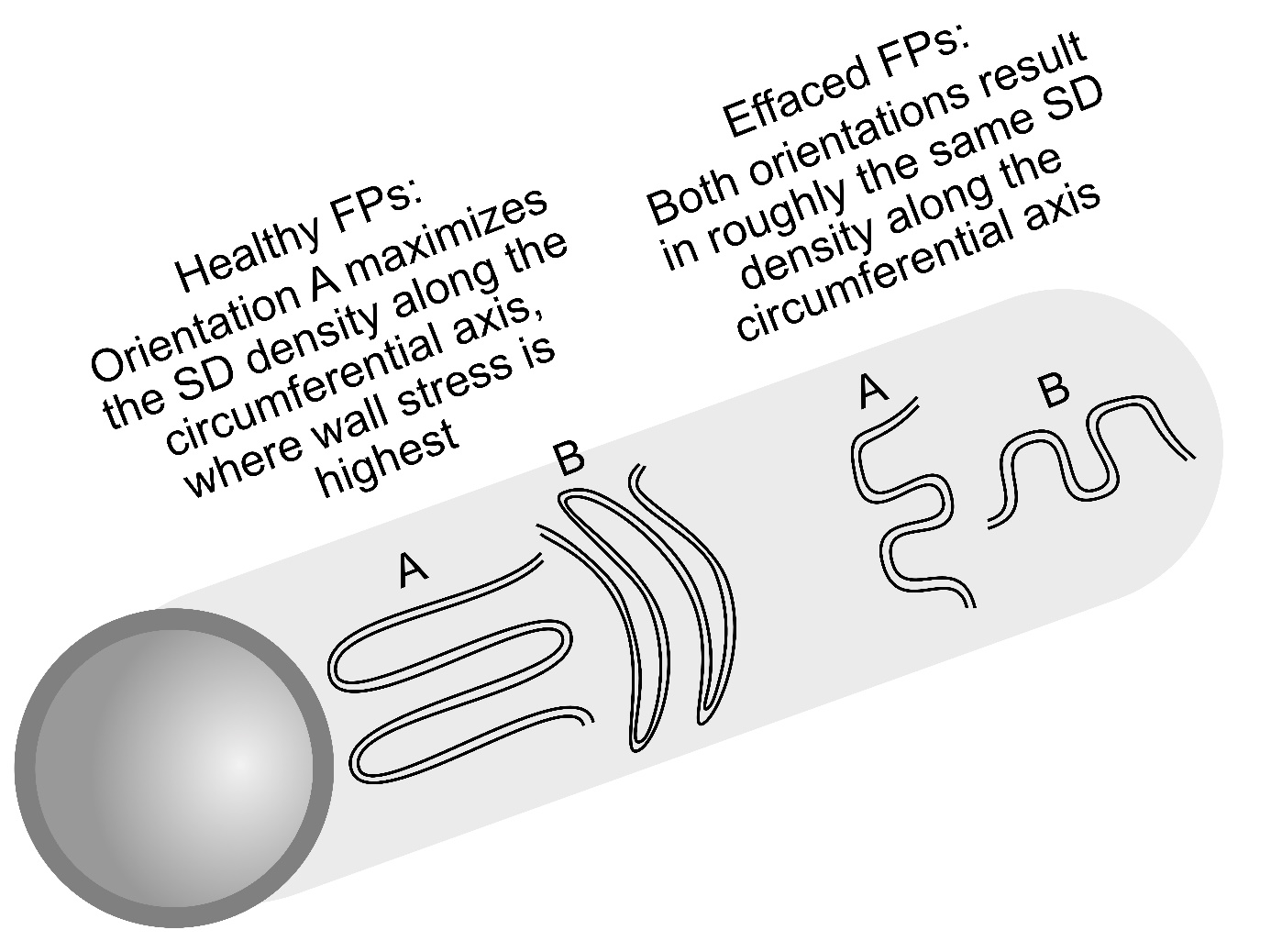

**Supplementary Figure 2.** Orientation of FPs is optimized under healthy conditions but not under pathological conditions. For healthy and elongated FPs, the observed orientation is parallel with the central axis of capillaries resulting in SD molecules being aligned in parallel with the most prominent circumferential wall stress component. For effaced FPs which takes on a shorter, wider and more circular shape, both orientations result in approximately the same number of SD molecules aligned in parallel with the circumferential wall stress component, possibly explaining the lack of preferred orientation in mice with glomerular disease.
